# A uniform *in vitro* efficacy dataset to guide antimicrobial peptide design

**DOI:** 10.1101/463588

**Authors:** Deepesh Nagarajan, Tushar Nagarajan, Neha Nanajkar, Nagasuma Chandra

## Abstract

Antimicrobial peptides are ubiquitous molecules that form the innate immune system of organisms across all kingdoms of life. Despite their prevalence and early origins, they continue to remain potent natural antimicrobial agents. Antimicrobial peptides are therefore promising drug candidates in the face of overwhelming multi-drug resistance to conventional antibiotics. Over the past few decades, thousands of antimicrobial peptides have been characterized *in vitro*, and their efficacy data is now available in a multitude of public databases. Computational antimicrobial peptide design attempts typically use such data. However, utilizing heterogenous data aggregated from different sources presents significant drawbacks. In this report, we present a uniform dataset containing 20 antimicrobial peptides assayed against 30 organisms spanning gram positive, gram negative, fungal, and mycobacterial origin. We draw inferences from the results of 600 individual MIC assays, and discuss what characteristics are essential for antimicrobial peptide efficacy. We expect our uniform dataset to be useful for future projects involving computational antimicrobial peptide design.

## 1 Introduction

Antimicrobial peptides (AMPs) form part of the armory used by the innate immune system of a variety of organisms ranging from microbes to humans. Despite their prevalence and early origins, they continue to remain potent natural antimicrobial agents. AMPs are therefore promising drug candidates^1^ in the face of overwhelming multi-drug resistance to conventional antibiotics^2^.

Most antimicrobial peptides are short molecules, ranging from 6-50 residues^3^. They are typically amphiphilic with a net positive charge^4^, although neutral^5^ and negatively charged peptides^6^ are also encountered. The primary mechanism of action of AMPs involves direct interaction with, and disruption of, the bacterial membrane. Positively charged antimicrobial peptides are attracted towards negatively charged phospholipid moieties, which facilitates AMP incorporation into the lipid bilayer. Post-incorporation, three models compete to explain AMP-induced membrane disruption: the toroidal-pore model^7^, the barrel stave model^8^, and the carpet model^9^. Although the mechanisms described in these models differ, all describe direct peptide incorporation into, and disruption of, bacterial membranes, leading to death.

Secondary mechanisms of action for AMPs have also been proposed, which include inhibition of aerobic electron transport^10^, inhibition of nucleotide^11, 12^ / protein^13^ synthesis, promotion of ribosomal aggregation,^14^, membrane protein delocalization^15^, and metabolic inhibition^14, 16^. Adding a further layer of complexity, many natural antimicrobial peptides possess weak bactericidal activity. Rather than directly inhibit bacterial growth, they are now known to act in concert with the host immune system through mechanisms including chemokine induction^17^, histamine release^18^, and angiogenesis modulation^19^. These immunomodulatory effects have only recently begun to receive attention.

Despite the complexities involved in understanding the mechanisms of action, several attempts at creating AMPs using rational design approaches have been made. Pexiganan^20^, for example, is a rationally designed Magainin-2 derivative that displays superior bactericidal properties. Other design approaches have involved employing simple sequence repeats that mimic the biophysical features of natural antimicrobial peptides. Leu-lys repeats^21^, trp-arg repeats^22^, and trp-leu-lys repeats^23^, have all displayed broad spectrum antimicrobial activity. A later study using more elaborate repeat patterns yielded similar results^24^. Computational approaches to AMP design have employed genetic algorithms^25^, QSAR approaches^26^, linguistic models^27^, and LSTM neural networks^10^.

A better understanding of the sequence and structural characteristics responsible for AMP activity would not only help to further understand the mechanisms of natural AMPs, but also form the basis for the *de novo* design of new AMPs. Essential to understanding these features is the availability of large datasets containing information on the efficacy of existing AMPs. Several databases curating thousands of antimicrobial peptides exist, such as the Antimicrobial Peptide Database (APD)^3^, Yet Another Database of Antimicrobial Peptides (YADAMP)^28^, the Collection of Antimicrobial Peptides (CAMP)^29^, and Data Repository of Antimicrobial Peptides (DRAMP)^30^. In all cases, minimum inhibitory concentration (MIC) data from different sources are compiled to form a single database. This approach is entirely reasonable given the heterogeneous nature of efficacy data available, but nevertheless suffers from significant drawbacks:

1. Individual studies report MIC values obtained using varying protocols, which produce different results.
2. Different groups use different type cultures of the same organism for MIC estimation.
3. Negative data (MIC results for ineffective peptides) is seldom published.

Therefore, MIC values obtained from different sources, but compiled within the same dataset, cannot directly be compared. Further, the lack of negative data limits computational design approaches that require diverse samples for training.

In this study, we report the MIC results of 20 antimicrobial peptides possessing diverse sequences, and possessing varying efficacy against 30 organisms spanning gram positive, gram negative, fungal, and mycobacterial origin. We report 600 individual MIC assays. While this data is quantitatively inferior to existing AMP databases (that contain thousands of MIC values), it is qualitatively superior. All MIC experiments were performed on the same strains for every organism, performed using the same protocol, and performed in the same laboratory by the same personnel, ensuring uniformity across the dataset and allowing direct intra-dataset comparisons to be made. Further, a preliminary analysis revealed sequential and structural traits responsible for AMP efficacy, proving the utility of our dataset for future AMP design projects.

## 2 Results

### 2.1 Peptides and cultures assayed

We synthesized and experimentally characterized 20 peptides. 10 sequences (NN2_0018 → NN2_0055) posed good antimicro-bial activity and were described in our previous work^10^. Another 10 sequences (NN2_0000 → NN2_0009) possessed poor antimicrobial activity. Although these peptides are mostly ineffective and possess no therapeutic potential, they can still be used to understand the sequence and structural characteristics of effective peptides. Sequences for all 20 peptides are provided in Table 1. All 20 peptides were readily soluble in distilled water at concentrations of ≥2 mg/ml. A broth microdilution method developed for cationic antimicrobial peptides^31^ was used for MIC determination. For MIC assays, peptide concentrations of 0.25*μ*g/ml → 128*μ*g/ml were used. Based on their diversity and clinical relevance, we selected 30 cultures were chosen for MIC testing. The cultures tested included Gram positive, Gram negative, fungal, and mycobacterial organisms. Wherever possible, cultures isolated from clinical samples were used. Most of our cultures were obtained from the Microbial Type Culture Collection (MTCC, Chandigarh, India). ATCC strains and reference strains were also used. Minimum inhibitory concentrations for all peptides and all cultures is provided in Table 2.

**Table 1.**
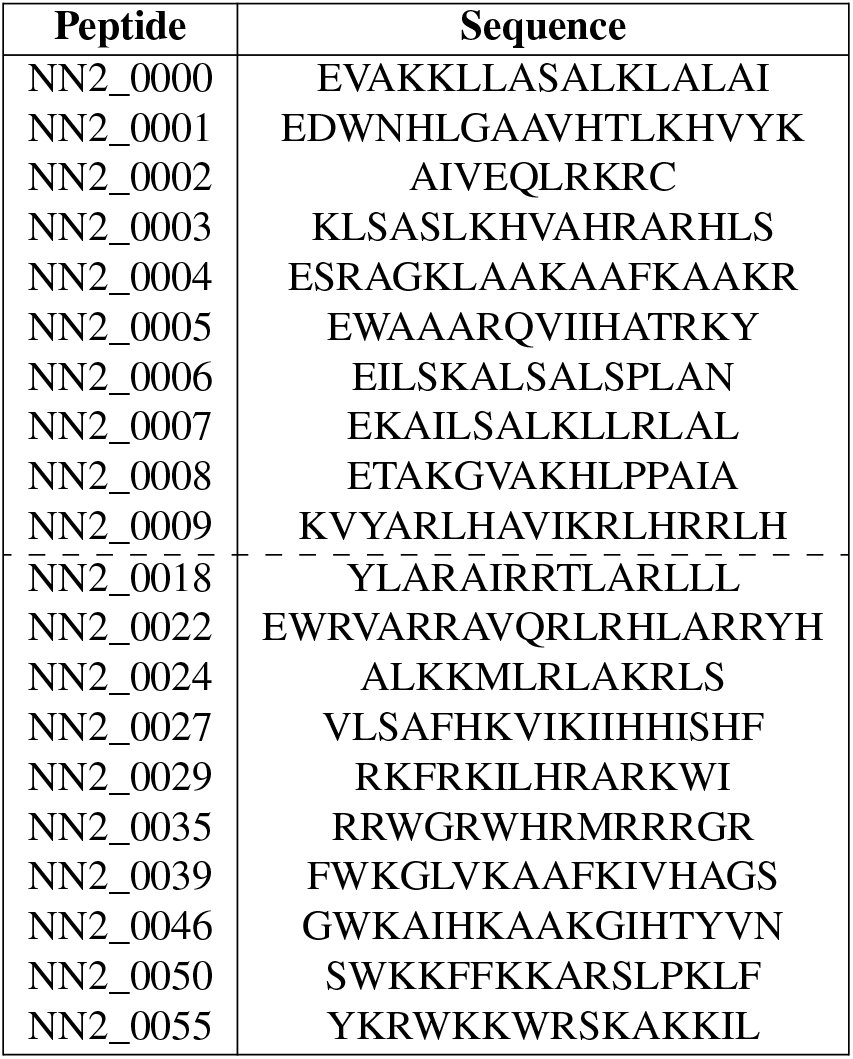
Names and sequences are provided for all peptides described in this study. Peptides NN2_0000 NN2_0009 are reported for the first time in this study. Peptides NN2_0018 NN2_0050 were reported and characterized in our previous work^10^.

**Table 2.** MIC values are provided in *μ*g/ml for all 20 peptides tested against 30 cultures. MIC values in **bold** are the lowest MIC values for a given culture. Cultures with MIC values exceeding 128 *μ*g/ml are reported as blank cells. *Peptide scores* are provided for all effective peptides. Culture names marked with an *asterisk* indicate that the resazurin protocol was used to estimate minimum bactericidal concentration (MBC). Note that MIC values for peptides NN2_0018 → NN2_0050 were reported in our previous work^10^. Cultures with an asterisk appended to their names displayed mucoid/plaque morphologies, and the MBC was determined using the resazurin method.

| Organism | Culture ID | NN2_0000 | NN2_0001 | NN2_0002 | NN2_0003 | NN2_0004 | NN2_0005 | NN2_0006 | NN2_0007 | NN2_0008 | NN2_0009 | NN2_0018 | NN2_0022 | NN2_0024 | NN2_0027 | NN2_0029 | NN2_0035 | NN2_0039 | NN2_0046 | NN2_0050 | NN2_0055 |
| --- | --- | --- | --- | --- | --- | --- | --- | --- | --- | --- | --- | --- | --- | --- | --- | --- | --- | --- | --- | --- | --- |
| *E. coli* | K12 MG1655 |  |  |  |  |  |  |  |  |  | 16 | 32 | 16 | 64 | 32 | 8 |  | 64 |  | **4** | **4** |
| *A. baumannii* | MTCC 9829 |  |  |  |  |  |  |  | 128 |  | 32 | 16 | 16 | 128 | 16 | 8 |  | 16 |  | **4** | 16 |
| *S. boydii* | MTCC 11947 | 128 |  |  |  |  |  |  | 128 |  | 8 | 32 | 8 | 64 | 16 | **1** | 64 |  |  | 2 | 8 |
| *S. flexnerii* | MTCC 1457 | 128 |  |  |  |  |  |  | 128 |  | 4 | 8 | **1** | 32 | 8 | 4 | 64 |  |  | 4 | 8 |
| *S. typhimurium* | ATCC 14028 |  |  |  |  |  |  |  |  |  | 32 | 32 | 16 |  |  | 16 |  |  |  | **8** | 32 |
| *S. enterica* | MTCC 9844 |  |  |  |  |  |  |  | 128 |  | 32 | 16 | 16 | 32 | 32 | 16 |  |  |  | **8** | 16 |
| *K. pneumoniae* | MTCC 7407 |  |  |  |  |  |  |  |  |  |  | **32** | 128 |  |  | 64 |  |  |  | **32** | 128 |
| *K. oxytoca* | MTCC 2275 |  | 128 |  |  |  |  |  |  |  | **8** | 16 | **8** | 128 |  | 32 | 64 | 64 |  | 16 | 32 |
| *P. aeruginosa* | MTCC 3542 |  |  |  |  |  |  |  |  |  |  |  | 128 |  |  | **16** |  |  |  | 32 | 128 |
| *P. vulgaris* | MTCC 1771 |  |  |  |  |  |  |  |  |  |  | 128 | 128 |  |  | 64 |  |  |  | **16** | 64 |
| *P. mirabilis* | MTCC 3158 |  |  |  |  |  |  |  |  |  |  |  |  |  |  |  |  |  |  |  |  |
| *C. koserii* | MTCC 1657 | 128 |  |  |  |  |  |  | 128 |  | 32 | 16 | 16 | 64 | 16 | 64 | 16 | 64 |  | **8** | 16 |
| *C. freundii* | MTCC 1658 |  |  |  |  |  |  |  |  |  | 32 | **16** | 64 |  |  | 32 |  |  |  | 32 | 128 |
| *N. mucosa* * | MTCC 1772 |  |  |  |  |  | 128 |  | 128 |  | 32 | **16** | 32 | 128 | 32 | 32 | 64 | 128 |  | **16** | 64 |
| *V. cholerae* | MTCC 3904 |  |  |  |  |  |  |  |  |  | 128 | **64** | 128 |  |  | 128 |  |  | 128 | **64** | 128 |
| *E. gergoviae* | MTCC 3826 |  |  |  |  |  |  |  |  |  | 128 | 128 | 128 |  |  |  |  |  |  | **64** |  |
| *H. influenzae* | MTCC 621 | 64 |  |  |  | 128 | 128 |  | 16 |  | 4 | 8 | 8 | 64 | 8 | 8 | 128 |  | 64 | **2** | 32 |
| *A. fecalis* | MTCC 1937 |  |  |  |  |  |  |  |  |  |  | **32** | 128 |  |  | 128 |  |  |  | 64 |  |
| *B. bronchiseptica* | MTCC 6837 | 16 |  |  |  |  |  |  | 64 |  | 4 | 4 | 2 | 8 | 4 | **1** |  | 8 | 128 | **1** | 8 |
| *E. aerogenes* | MTCC 111 |  |  |  |  |  |  |  |  |  |  | 32 |  |  |  |  |  |  |  | **16** | 128 |
| *S. maltophilia* | MTCC 1890 | 128 |  |  |  |  |  |  |  |  |  | **16** | 64 |  |  | 128 |  | 128 |  | **16** | 128 |
| *M. luteus* * | MTCC 425 | 32 |  | 128 | 32 | 32 | 32 |  | 0.5 |  | 2 | 2 | 2 | 8 | 1 | **0.25** | **0.25** | 2 | 64 | 2 | 0.5 |
| *S. aureus* | MTCC 3160 |  |  |  |  |  |  |  |  |  |  | **16** |  |  |  | 128 |  |  |  | 128 |  |
| *S. hemolyticus* | MTCC 3383 |  |  |  |  |  |  |  | 128 |  | **4** | 16 | 16 | 128 | 32 | 8 | **4** | 16 |  | 8 | **4** |
| *E. faecalis* | MTCC 439 |  |  |  |  |  |  |  |  |  |  | **64** |  |  |  |  |  |  |  | 128 |  |
| *C. glutamicum* | MTCC 2679 | 32 |  |  |  |  | 64 |  | 32 |  | 2 | 4 | **1** | 16 | 4 | 2 | 4 | 2 | 64 | 2 | 2 |
| *C. pseudoTB* * | MTCC 3158 |  |  |  |  |  |  |  |  |  |  | **128** |  |  |  |  |  |  |  |  |  |
| *B. alcalophilis* | MTCC 860 |  |  |  |  |  |  |  | 64 |  | 32 | **16** | **16** |  | 32 | 64 | 32 | **16** |  | 32 | 64 |
| *M. smegmatis* * | MC2155 |  |  |  | 128 |  |  |  |  |  | 32 | 64 | **16** | 64 | 32 | **16** | **16** | 128 |  | 64 | 32 |
| *C. albicans* * | MTCC 425 |  |  |  |  |  |  |  |  |  | **32** | 128 | 64 | 128 |  | 64 | 64 |  |  | 64 | 64 |
| net charge |  | 3 | 2 | 3 | 4 | 6 | 3 | 1 | 3 | 2 | 6 | 4 | 7 | 5 | 2 | 7 | 8 | 3 | 3 | 6 | 8 |
| peptide score |  |  |  |  |  |  |  |  |  |  | 3 | 10 | 5 |  |  | 6 | 3 | 1 |  | 14 | 2 |

### 2.2 Identifying effective peptides based on MIC data

We identifed effective, broad-spectrum peptides, using a relative scoring scheme^10^. Simply described, for a given peptide, its *peptide score* was calculated by counting number of cultures it inhibited with the lowest MIC(in comparison to the MICs of all other peptides for a given culture). A mathematical description of the peptide score is provided in Equation 1.

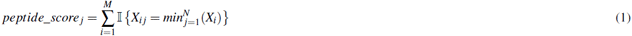

For this equation:

- *X*: a matrix of MIC values.
- *M*: rows containing MIC values for a given organism.
- *N*: columns containing MIC values for a given peptide.
- 0 ≤ *i* ≤ *M*, 0 ≤ *j* ≤ *N*
- Multiple minimum MIC values can occur along a given row.

### 2.3 MIC experiments suggest a common mechanism of action for both Gram positive and Gram negative organisms

From Table 2, it is apparent that peptides displaying a broad spectrum of activity also inhibit cultures at lower concentrations (low MIC values). Conversely, peptides displaying a narrower spectrum of activity inhibit cultures at higher concentrations (high MIC values). These trends are illustrated in Figure 1A, and were observed to be strongly correlated (r = −0.83).

**Figure 1.**
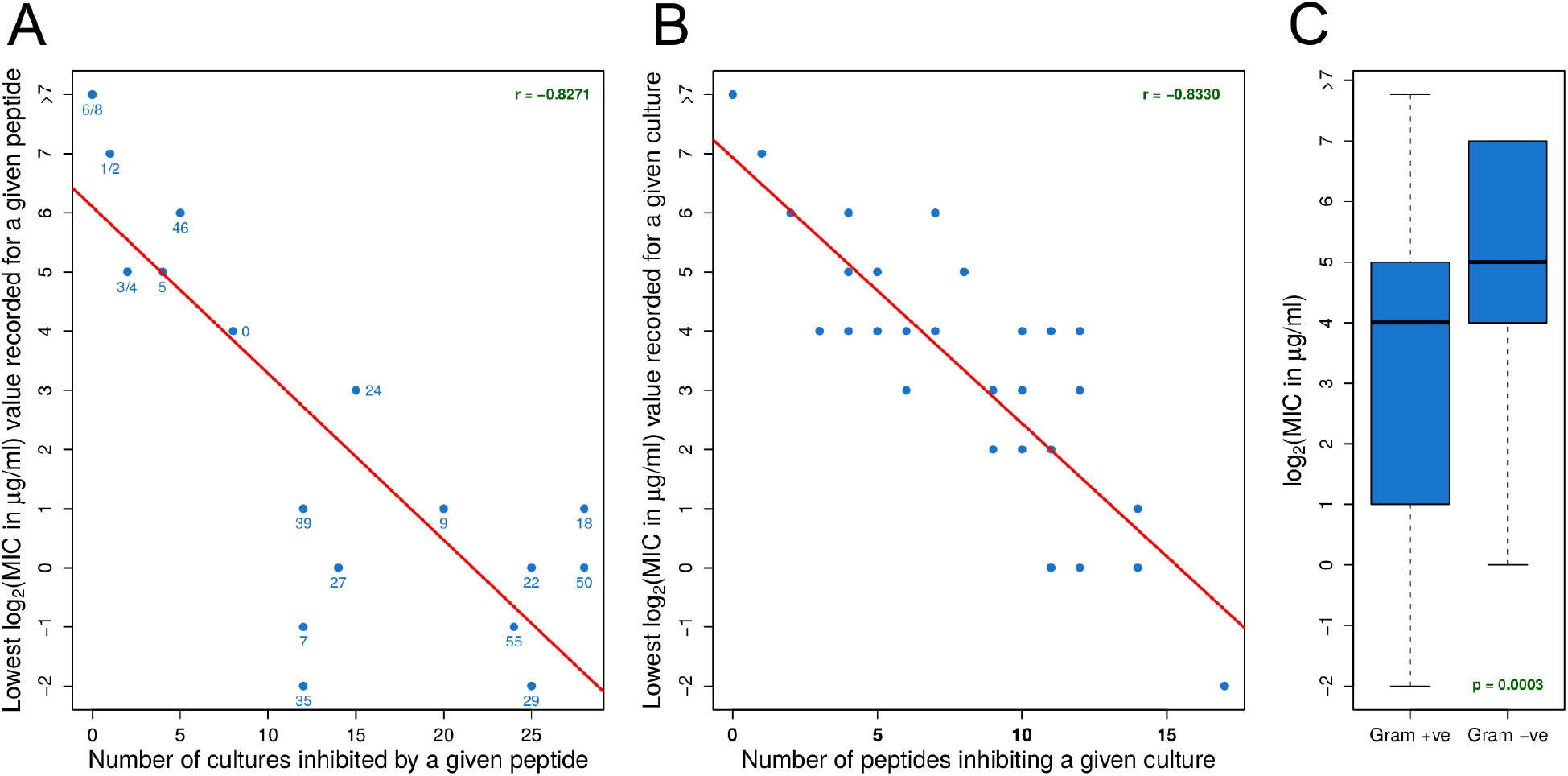
Understanding how susceptibility, spectrum of activity, and Gram nature relate to a common mechanism of action. **(A)** Peptide-oriented scatterplot displaying the strong correlation observed between the lowest MIC value recorded for a given culture, and the number of cultures inhibited. The numbering represents individual peptides (eg: NN2_0050 → 50). **(B)** Culture-oriented scatterplot displaying the strong correlation observed between lowest MIC value recorded for a given peptide, and the number of peptides inhibiting the given culture. **(C)** Boxplots depicting the small but statistically significant difference between MIC values obtained for all Gram positive organisms, as compared to those obtained for all Gram negative organisms. The Welch 2-sample t-test was used to determine statistical significance.

These trends were mirrored from the perspective of the cultures tested. Cultures inhibited at lower concentrations (low MIC values) by any peptide were found to be inhibited by a larger number of peptides. Conversely, cultures inhibited at higher concentrations (high MIC values) by any peptide were found to be inhibited by fewer peptides. Once again, as illustrated in Figure 1B, these variables were observed to be strongly correlated (r = −0.83).

From these strongly correlated observations, two inferences can be made:

1. For an organism, susceptibility to one effective peptide indicates greater susceptibility to all effective peptides.
2. For an effective peptide, efficacy on one organism indicates greater efficacy on all organisms.

This indicates that all the peptides found to be effective possess very similar mechanisms of action, despite differences in their size and sequence. Further, this mechanism is conserved across diverse organisms. Therefore, these peptides would only differ quantitatively in their degree of efficacy while following the same qualitative mechanism of action.

All peptides were found to inhibit both Gram positive and Gram negative cultures. However, we observed a small but statistically significant difference in the susceptibility of Gram positive organisms as compared to their Gram negative counterparts (Figure 1C). Ignoring susceptibilities >128 *μ*g/ml, the median MIC of Gram positive organisms for all peptides tested was 16 *μ*g/ml, 2-fold lower than the corresponding Gram negative median MIC of 32 *μ*g/ml (p = 0.0024). These observations remained statistically significant even after including susceptibilities >128 *μ*g/ml (p = 0.0035). Since no peptides were observed to display selective activity against either Gram positive or Gram negative cultures, these observations are once again best explained by a similar mechanism of action. Gram positive organisms may be inherently more susceptible to antimicrobial peptides. Therefore, peptides would act with a similar mechanism in Gram positive organisms, differing only in the magnitude of inhibition compared to their Gram negative counterparts.

### 2.4 Positively charged residues are associated with increased peptide activity

Trends between peptide positive charge, apolar content, and antimicrobial activity are illustrated in Figure 2. From this figure, it is clear that peptides possessing a low per-residue positive charge of ≤ +0.1 are ineffective (100% of all MIC values were > 128 *μ*g/ml) (Figure 2A). However, peptides possessing a high per-residue positive charge of +0.5 → +0.6 display submicromolar MIC values. These results are expected, as cationic antimicrobial peptides are a well-established family of AMPs. For these peptides, positively charged residues allow it to interact with, and disrupt, the negatively charged bacterial membrane. Statistical significance was calculated by dividing the data at the median positive charge per residue (0.25). The difference in MIC distributions between the low-positive and high-positively charged peptide datasets was statistically significant (p = 2.2e-16, Fisher’s test).

**Figure 2.**
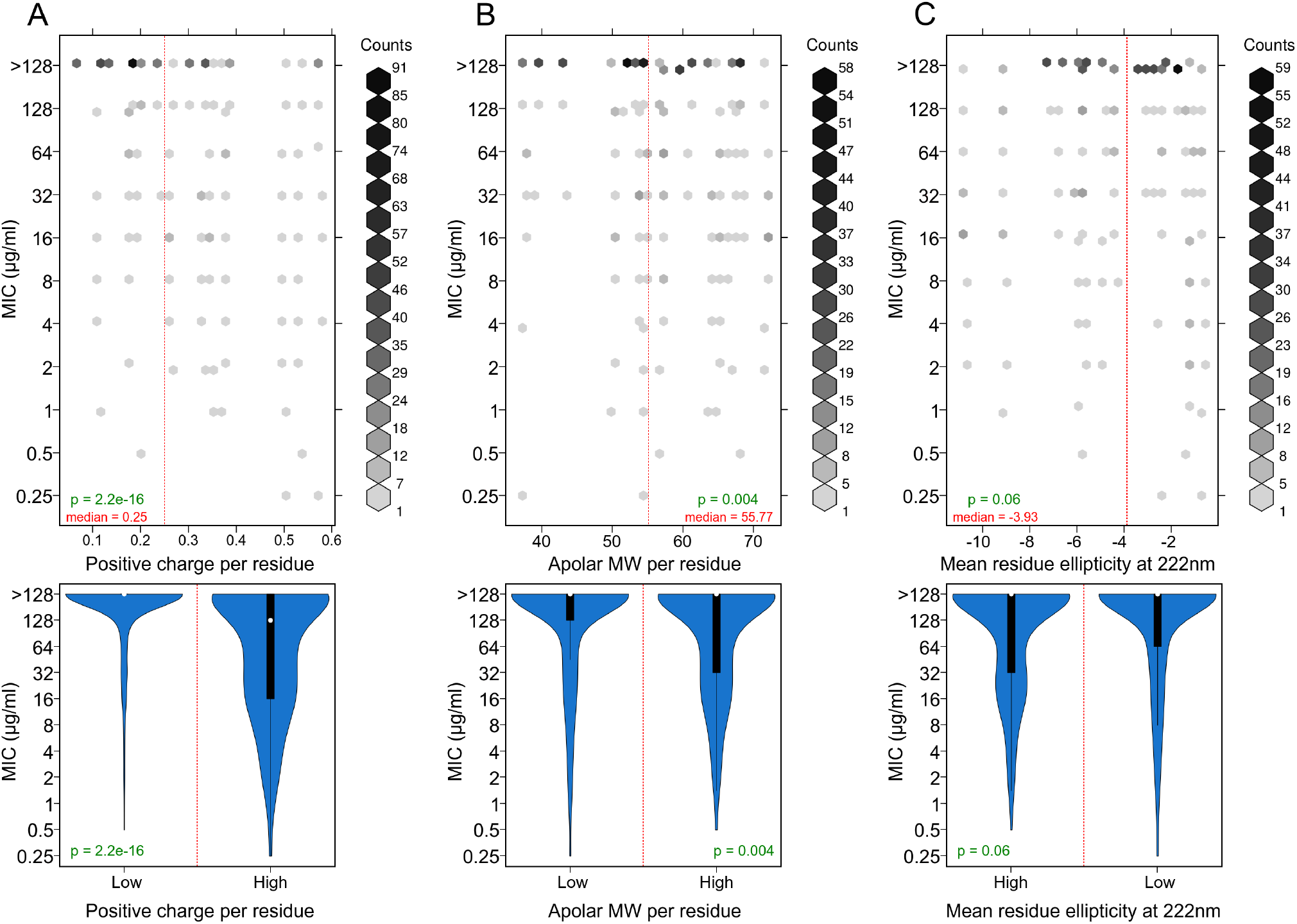
Trends between peptide positive charge, apolar content, helicity, and antimicrobial activity are illustrated as 2D histograms **(above)** and violinplots **(below)**. From 20 peptides x 30 cultures, 600 individual MIC experiments were performed. **(A)** Peptides possesing higher per-residue positive charges also possess lower MIC values and greater effiacy. Peptide positive charge per residue is merely the sum of (Lys +Arg-Asn-Glu residues)/peptide length. **(B)** Peptides possessing higher per-residue apolar content also possess lower MIC values and greater effiacy. Apolar content is merely the total molecular weight of all apolar residues (AVLIMFYWPC) / peptide length. **(C)** Peptide secondary structure content, as determined using circular dichroism, is not linked to MIC values or peptide efficacy. Note that mean residue ellipticity is measured in 10^3^ deg cm^2^ dmol^−1^ units.

### 2.5 Apolar residues are associated with increased peptide activity

Peptides possessing greater apolar molecular weights per residue displayed slightly lower MIC values, and therefore slightly greater efficacy (Figure 2B). Statistical significance was calculated by dividing the data at the median apolar molecular weight per per residue (55.77). The difference in MIC distributions between the relatively polar and apolar peptide datasets was statistically significant (p = 0.004, Fisher’s test). These results indicate that designed peptides would benefit from the inclusion of large apolar residues such as Phe, Tyr, and Trp in their sequence. It should be noted that this trend appeared much weaker than for peptide positive charge.

### 2.6 Helicity is not essential for peptide activity

Circular dichroism (CD) experiments were performed to investigate the secondary structural characteristics of all designed peptides. Near-UV CD experiments in trifluoroethanol (TFE) were performed to determine peptide helical content. Trifluoroethanol is a low-dielectric solvent that encourages helix formation. It is used to mimic the bacterial membrane environment^32, 33^ while investigating the properties of antimicrobial peptides.

In an aqueous solution, all peptide designs adopted the random coil conformation, displaying a characteristic minima beyond 200 nm (195 nm). However, upon increasing the concentration of trifluoroethanol, some peptides underwent conformational changes, adopting alpha helical structures. In a solution of 40% trifluoroethanol, 11 of the 20 designed peptides displayed some degree of alpha helicity. These peptides displayed a characteristic alpha-helical double minima at 208 nm and 222 nm. The 11 helical peptides are as follows: NN2_0000, NN2_0001, NN2_0004, NN2_0005, NN2_0007, NN2_0009, NN2_0018, NN2_0022, NN2_0024, NN2_0027, and NN2_0039. The other designed peptides adopted random coil conformations, even upon addition of 40% trifluoroethanol. These results are depicted in Figure 3.

**Figure 3.**
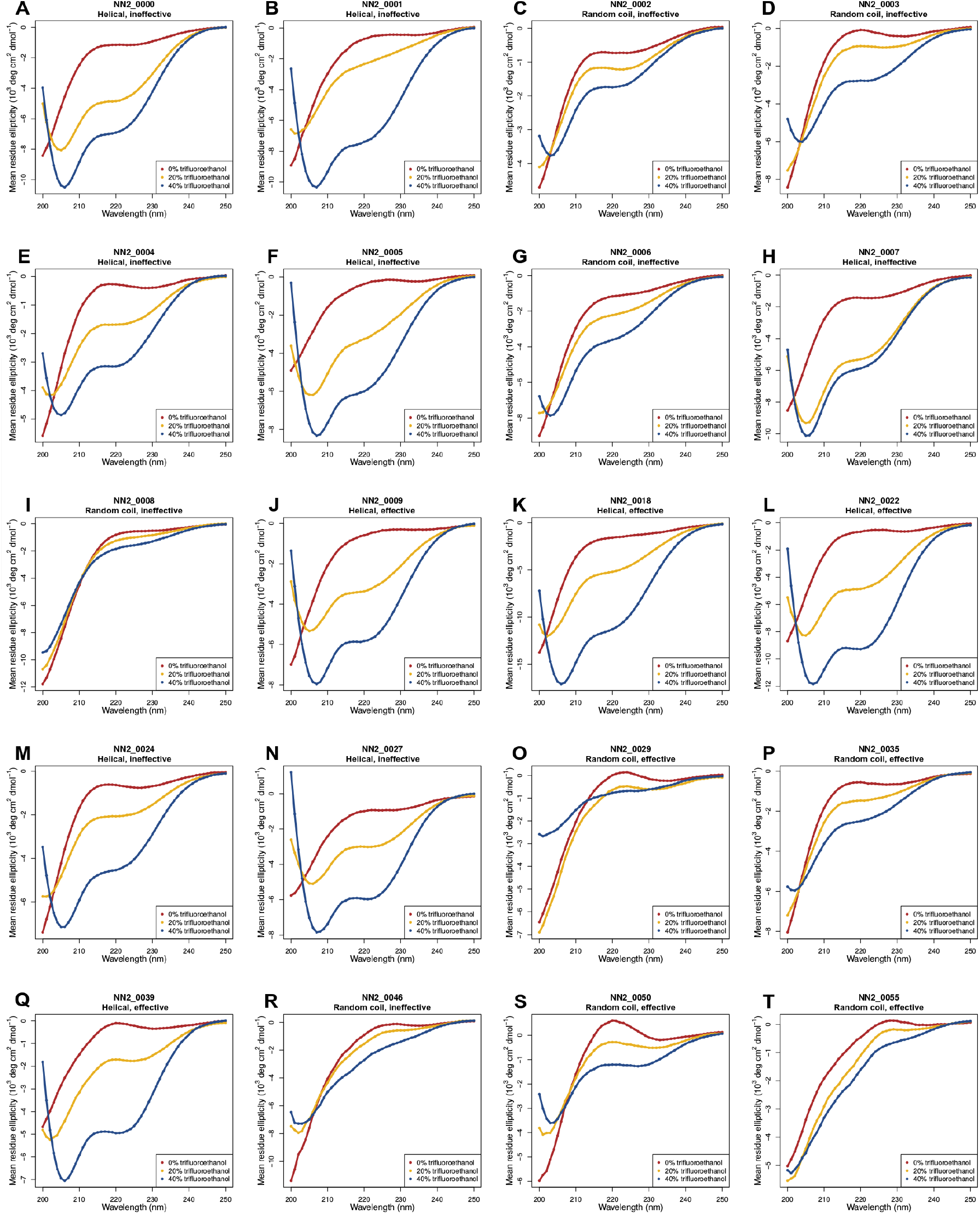
Circular dichroism (CD) experiments. **(A-T)** The far-UV circular dichroism spectra of all 20 designed peptides was collected. Each peptide was dissolved in aqueous solvent containing 0% trifluoroethanol (in red), 20% trifluoroethanol (in yellow), and 40% trifluoroethanol (in blue), in order to study the secondary structures adopted during peptide-membrane interaction. The secondary structure and relative efficacy of each peptide are provided below the title of each graph. Note that "effective" and "ineffective" are relative terms based on the peptide score, as described in the text. A peptide labeled as "ineffective" may still display antimicrobial activity against various cultures, but only to a lesser extent as compared to peptides labeled as "effective".

From the CD spectra observed, it is apparent that alpha helicity was not an essential factor for antimicrobial activity Figure 2C. Statistical significance was calculated by dividing the data at the median *mean residue ellipticity* (−3.93) measured at 222 nm, and measured in 40% trifluoroethanol. The difference in MIC distributions between the helical and non-helical peptide datasets was not statistically significant (p = 0.06, Fisher’s test).

## 3 Discussion

In this work, we present a dataset containing 600 MIC values obtained from testing 20 peptides against 30 diverse pathogens. Gram positive, gram negative, fungal, and mycobacterial isolates were tested. As our data was generated using the same protocol^31^, our peptides were tested against the same type culture for every organism, and we have included negative data in the form of ineffective peptides (NN2_0000 → NN2_0008). Therefore, our data is qualitatively superior to aggregated, multi-source heterogeneous data found on antimicrobial peptide databases^3, 28–30^, and should therefore be more suitable for training future AMP design algorithms.

We have also performed simple statistical analyses of our data, and determined that positive charge is essential for AMP efficacy (Figure 2A). Further, a large apolar residue content also contributes to AMP efficacy Figure 2B). These results agree with previously understood mechanisms of AMP action^14^. Indeed *de novo* peptides possessing trp-arg repeats^22^ and trp-leu-lys repeats^23^ were designed utilize the same principles.

Counter-intuitively, we observed that alpha helicity was not required for peptide efficacy (Figures 2C, 3). However, this result can be explained the carpet model AMP activity. Briefly, positively charged amphiphilic peptides, with either monomeric or random structures, are described to cover the cell membrane in a *carpet-like* manner. Once a threshold concentration is reached, the peptides disrupt the bilayer curvature, disintegrating the membrane. The competing toroidal pore^7^ and barrel stave^8^ models describe the insertion of alpha helical peptides perpendicular to the cell membrane, forming nanometer-scale pores that lead to the leakage of cellular contents and ultimately death. These observations favor the carpet model:

1. Peptides adopting both alpha helical and random coil structures were found to be effective antimicrobial agents (Figures 2C, 3). Random coils cannot form the nanometer-scale pores described by the toroidal pore and barrel stave models.
2. Our previous work^10^ reported prominent blebbing observed on the *S. haemolyticus* cell membrane, and large-scale membrane damage observed on *E. coli*, upon treatment with peptides NN2_0018 and NN2_0050. These disruptions cannot be explained through the formation of nanometer-scale pores alone. Previously, the carpet model had successfully explained similar blebbing on the *P. aeruginosa* cell membrane^9^.

Ultimately, the main contribution of this work is the homogeneous AMPs dataset, which should provide valuable training data for the design of new AMPs. New drugs of all classes are urgently needed to combat the emergence of multidrug resistant pathogens.

## 4 Methods

### 4.1 Peptide synthesis

GenScript, Inc. supplied all the peptides used in this study. 20 mg of the 20 NN2-family peptides were synthesized by GenScript as part of a peptide library.

### 4.2 Antimicrobial susceptibility assays

The microwell dilution method as described by Wiegand et. al.^31^ (Protocol E: Broth microdilution for antimicrobial peptides that do not require the presence of acetic acid/BSA). This protocol was especially optimized for the MIC determination of cationic antimicrobial peptides, and involves the use of polypropylene rather than polystyrene 96-well plates.

In order to estimate the MICs of cultures displaying plaque or mucoid morphologies, we used a modified protocol involving resazurin. Resazurin is normally a marginally fluorescent dye. However, microbial aerobic respiration reduces it to the highly fluorescent resorufin form. After incubating microbial cultures at 37 °C for 12 hours (according to protocol E), 30 *μ*l of a 0.02% (w̌) aqueous resazurin solution was pipetted into each well of a 96-well polypropylene plate. Further incubation at 37 °C for 12 hours was followed by fluorescence detection (excitation: 530 nm, emission: 590 nm) to determine cell viability. Since bacterial respiration is a measure of cell viability, this mechod calculates minimum bactericidal concentrations(MBCs) instead of minimum inhibitory concentrations (MICs).

### 4.3 Circular dichroism experiments

All circular dichroism (CD) experiments were performed using the Jasco J-810 spectrophotometer. A 1mm path-length quartz cuvette with a sample volume of 300 *μ*l was used. Far-ultraviolet spectra (200-250 nm) were collected at a 3 nm bandwidth and with a 4 s response-time. Every spectrum was collected in triplicate and averaged. Buffer spectrum correction was also performed.

CD experiments were performed to understand the changes in antimicrobial peptide secondary structure during peptide-membrane interaction. Trifluoroethanol was chosen as a membrane mimic. Trifluoroethanol acts as both an apolar solvent, and as an agent to encourage helix formation. Trifluoroethanol-water solutions containing 0%, 20%, and 40% trifluoroethanol were prepared and used for all experiments.

## 5 Acknowledgements

The authors thank the Department of Biotechnology (DBT) and the Department of Science and Technology (DST), India, for funding this work.

## 6 Author contributions statement

D.N. Performed all MIC experiments (Table 2), circular dichroism experiments (Figure 3), and statistical analyses (Figure 2). T.N. designed the antimicrobial peptide sequences used in this study (Table 1). N.N. analyzed and interpreted trends in the MIC data (Figures 1). D.N. and N.C. conceived and designed all experiments. N. C. coordinated the study, planned experiments, and provided resources. All authors reviewed the results and approved the final version of the manuscript.

## References

1. Fox, J. L. Antimicrobial peptides stage a comeback. Nat. biotechnology 31, 379–382 (2013).

2. Vincent, J.-L. et al. Assessment of the worldwide burden of critical illness: the intensive care over nations (icon) audit. The lancet Respir. medicine 2, 380–386 (2014).

3. Wang, Z. & Wang, G. Apd: the antimicrobial peptide database. Nucleic acids research 32, D590–D592 (2004).

4. Scott, M. G. & Hancock, R. E. Cationic antimicrobial peptides and their multifunctional role in the immune system. Critical Rev. Immunol. 20 (2000).

5. Hotchkiss, R. D. & Dubos, R. J. Bactericidal fractions from an aerobic sporulating bacillus. J. Biol. Chem. 136, 803–804 (1940).

6. Harris, F., Dennison, S. R. & Phoenix, D. A. Anionic antimicrobial peptides from eukaryotic organisms. Curr. Protein Pept. Sci. 10, 585–606 (2009).

7. Matsuzaki, K., Murase, O., Fujii, N. & Miyajima, K. An antimicrobial peptide, magainin 2, induced rapid flip-flop of phospholipids coupled with pore formation and peptide translocation. Biochemistry 35, 11361–11368 (1996).

8. Baumann, G. & Mueller, P. A molecular model of membrane excitability. J. supramolecular structure 2, 538–557 (1974).

9. Shai, Y. Mode of action of membrane active antimicrobial peptides. Pept. Sci. 66, 236–248 (2002).

10. Nagarajan, D. et al. Computational antimicrobial peptide design and evaluation against multidrug-resistant clinical isolates of bacteria. J. Biol. Chem. 293, 3492–3509 (2018).

11. Yonezawa, A., Kuwahara, J., Fujii, N. & Sugiura, Y. Binding of tachyplesin i to dna revealed by footprinting analysis: significant contribution of secondary structure to dna binding and implication for biological action. Biochemistry 31, 2998–3004 (1992).

12. Subbalakshmi, C. & Sitaram, N. Mechanism of antimicrobial action of indolicidin. FEMS microbiology letters 160, 91–96 (1998).

13. Patrzykat, A., Friedrich, C. L., Zhang, L., Mendoza, V. & Hancock, R. E. Sublethal concentrations of pleurocidin-derived antimicrobial peptides inhibit macromolecular synthesis in escherichia coli. Antimicrob. agents chemotherapy 46, 605–614 (2002).

14. Brogden, K. A. Antimicrobial peptides: pore formers or metabolic inhibitors in bacteria? Nat. Rev. Microbiol. 3, 238–250 (2005).

15. Wenzel, M. et al. Small cationic antimicrobial peptides delocalize peripheral membrane proteins. Proc. Natl. Acad. Sci. 111, e1409–E1418 (2014).

16. Gusman, H. et al. Salivary histatin 5 is an inhibitor of both host and bacterial enzymes implicated in periodontal disease. Infect. immunity 69, 1402–1408 (2001).

17. Nijnik, A. et al. Synthetic cationic peptide idr-1002 provides protection against bacterial infections through chemokine induction and enhanced leukocyte recruitment. The J. Immunol. 184, 2539–2550 (2010).

18. Befus, A. D. et al. Neutrophil defensins induce histamine secretion from mast cells: mechanisms of action. The J. Immunol. 163, 947–953 (1999).

19. Koczulla, R. et al. An angiogenic role for the human peptide antibiotic ll-37/hcap-18. The J. clinical investigation 111, 1665–1672 (2003).

20. Ge, Y. et al. In vitro antibacterial properties of pexiganan, an analog of magainin. Antimicrob. agents chemotherapy 43, 782–788 (1999).

21. Blondelle, S. E. & Houghten, R. A. Design of model amphipathic peptides having potent antimicrobial activities. Biochemistry 31, 12688–12694 (1992).

22. Deslouches, B. et al. Rational design of engineered cationic antimicrobial peptides consisting exclusively of arginine and tryptophan, and their activity against multidrug-resistant pathogens. Antimicrob. agents chemotherapy 57, 2511–2521 (2013).

23. Deslouches, B. et al. De novo generation of cationic antimicrobial peptides: influence of length and tryptophan substitution on antimicrobial activity. Antimicrob. agents chemotherapy 49, 316–322 (2005).

24. Wiradharma, N. et al. Synthetic cationic amphiphilic *α*-helical peptides as antimicrobial agents. Biomaterials 32, 2204–2212 (2011).

25. Fjell, C. D., Jenssen, H., Cheung, W. A., Hancock, R. E. & Cherkasov, A. Optimization of antibacterial peptides by genetic algorithms and cheminformatics. Chem. biology & drug design 77, 48–56 (2011).

26. Maccari, G. et al. Antimicrobial peptides design by evolutionary multiobjective optimization. PLoS Comput. Biol 9, e1003212 (2013).

27. Loose, C., Jensen, K., Rigoutsos, I. & Stephanopoulos, G. A linguistic model for the rational design of antimicrobial peptides. Nature 443, 867–869 (2006).

28. Piotto, S. P., Sessa, L., Concilio, S. & Iannelli, P. Yadamp: yet another database of antimicrobial peptides. Int. journal antimicrobial agents 39, 346–351 (2012).

29. Thomas, S., Karnik, S., Barai, R. S., Jayaraman, V. K. & Idicula-Thomas, S. Camp: a useful resource for research on antimicrobial peptides. Nucleic acids research gkp1021 (2009).

30. Fan, L. et al. Dramp: a comprehensive data repository of antimicrobial peptides. Sci. reports 6, 24482 (2016).

31. Wiegand, I., Hilpert, K. & Hancock, R. E. Agar and broth dilution methods to determine the minimal inhibitory concentration (mic) of antimicrobial substances. Nat. protocols 3, 163–175 (2008).

32. Gesell, J., Zasloff, M. & Opella, S. J. Two-dimensional 1h nmr experiments show that the 23-residue magainin antibiotic peptide is an *α*-helix in dodecylphosphocholine micelles, sodium dodecylsulfate micelles, and trifluoroethanol/water solution. J. biomolecular NMR 9, 127–135 (1997).

33. Conlon, J. M., Kolodziejek, J. & Nowotny, N. Antimicrobial peptides from ranid frogs: taxonomic and phylogenetic markers and a potential source of new therapeutic agents. Biochimica et Biophys. Acta (BBA)-Proteins Proteomics 1696, 1–14 (2004).

